# Dual RNA-seq in an aphid parasitoid reveals plastic and evolved adaptation to symbiont-conferred resistance

**DOI:** 10.1101/2019.12.13.875104

**Authors:** Alice B. Dennis, Heidi Käch, Christoph Vorburger

**Author notes:** Corresponding author: Alice Dennis, Unit of Evolutionary Biology/Systematic Zoology, Institute of Biochemistry and Biology, University of Potsdam, Karl-Liebknecht-Straße 24-25, 14476 Potsdam, Germany.

## Abstract

Coevolving taxa offer the opportunity to study the genetic basis of rapid reciprocal adaptation. We have used experimental evolution to examine adaptation in the parasitoid wasp *Lysiphlebus fabarum* to resistance conferred by the protective endosymbiont *Hamiltonella defensa* in its aphid host *Aphis fabae*. To examine a key stage in parasitoid infection, we have used RNA-seq to study gene expression in 4-5 day old parasitoid larvae contained in still living aphids. With this dual RNA-seq we can simultaneously view expression in individual experimentally evolved parasitoids and the aphids that house them. This gives a view of the sweeping changes in both taxa accompanying successful or unsuccessful infections. Among successful larvae, we find that experimentally evolved populations adapted to *H. defensa*-protected hosts differ in the expression of genes that include putative toxins and genes to cope with stress. These differences remain even when the larvae are developing in aphids possessing no defensive endosymbionts, suggesting that they are genetically based. In contrast, plastic responses between parasitoids reared in hosts with and without *H. defensa* are relatively small. Although aphids rely largely on their secondary endosymbionts for defense against parasitoids, we identify expression differences in aphids housing different parasitoid phenotypes. Together, these results demonstrate that wild parasitoid populations possess the genetic variation for rapid adaptation to host resistance, resulting in genetically based differences in gene expression that increase their success in parasitizing symbiont-protected host aphids.

## 1. Introduction

Genetic diversity is a fundamental requirement for evolutionary change. How diversity is created and maintained is an important and long-standing question in evolutionary biology. Some of the best examples of ongoing and rapid evolution come from host-parasite systems, where reciprocal selection between antagonists is particularly strong (Dheilly *et al.* 2015b; Thompson 1994; Windsor 1998). Host-parasite interactions can maintain rare and diverse phenotypes through negative frequency dependent selection, and drive diversification under certain conditions (Rouchet & Vorburger 2012; Woolhouse *et al.* 2002; Yoder & Nuismer 2010). Determining mechanisms of adaptation in host-parasite systems is thus important to understand their coevolution and how it drives specialization, and ultimately diversification.

Parasitoids are a special type of parasite that usually kill their host. Most described parasitoids are insects, and the majority of these are hymenoptera (Godfray 1994; Poirié *et al.* 2014). Parasitoid wasps that target aphids are a diverse group in themselves (Starý 1970), comprising highly specialized species exploiting just as single host species as well as more generalist taxa with a broader host range (Starý 2006; Straub *et al.* 2011). Successful infection of aphids requires an intricate oviposition behavior, during which female wasps also inject a complex venom arsenal, containing a breadth of rapidly evolving toxins that act to compromise aphids’ immunity (Colinet *et al.* 2014; Poirié *et al.* 2014) and redirect their resources from aphid reproduction to the developing parasitoid (Digilio *et al.* 2000).

Aphids are often protected from wasp infection by a suite of facultative heritable endosymbionts. These protective symbionts differ from their obligate symbionts, which provide key nutrients and share a long evolutionary history with their hosts (Bennett & Moran 2015). Facultative endosymbionts are not strictly required for host survival, but some provide protection from natural enemies, including parasitoid wasps (Oliver *et al.* 2003; Vorburger *et al.* 2010) and pathogenic fungi (Łukasik *et al.* 2013; Scarborough *et al.* 2005). One of the best-studied defensive symbionts is the γ-proteobacterium *Hamiltonella defensa* (Moran *et al.* 2005b), first discovered in pea aphids (*Acyrthosiphon pisum*) and shown to increase resistance to their parasitoid *Aphidius ervi* (Oliver *et al.* 2003). The ability to protect aphids against parasitoids has been linked to the presence of a toxin-encoding bacteriophage called APSE in *H. defensa*’s genome (Brandt *et al.* 2017; Oliver *et al.* 2009). Different strains of *H. defensa* may carry variants of APSE that encode different toxins, and this in turn correlates with different levels of protection provided by the symbiont strains (Moran *et al.* 2005a; Weldon & Oliver 2016). *Hamiltonella defensa* is widespread in aphids and is estimated to occur in >1,700 species (Oliver & Higashi 2018). In the black bean aphids (*Aphis fabae*) studied here, all *H. defensa* strains tested so far provide protection against this species’ most important parasitoid, *L. fabarum* (Cayetano *et al.* 2015; Vorburger *et al.* 2009). However, this protection shows remarkable specificity. Particular strains of *H. defensa* strongly increase resistance to some genotypes of *L. fabarum* but not others, and the spectrum of protection varies among *H. defensa* strains (Cayetano *et al.* 2015; Cayetano & Vorburger 2013; Schmid *et al.* 2012). The outcome of a parasitoid attack on a *H. defensa*-protected aphid is thus mediated by a genotype-by-genotype interaction between the parasitoid and the defensive symbiont.

We have recently employed experimental evolution to replicate populations of *L. fabarum*, allowing them to evolve on aphid hosts that were either uninfected with any defensive symbionts or infected with one of two different strains of *H. defensa*. These two strains carry different variants of the APSE phage that encode for different toxins (Figure 1, Dennis *et al.* 2017). As seen previously in parasitoids reared in protected host aphids (Dion *et al.* 2011; Rouchet & Vorburger 2014), we observed a rapid increase in parasitoid abilities to infect protected hosts. After only ten generations of experimental evolution, parasitoids evolving on *H. defensa*-protected aphids showed improved infectivity, suggesting that we have selected for standing genetic variation in the parasitoids’ ability to overcome symbiont-conferred resistance. Importantly, adaptation in experimental populations was specific to the *H. defensa* strain the parasitoids evolved with, and did not improve their infectivity in aphids protected by the other strain or in the unprotected (H-) control aphids. Comparisons of gene expression among adult female wasps from these lines found that this specificity was reflected in patterns of gene expression. In the treatments selected in the presence of *H. defensa*, most differentially expressed genes were specific to just that lineage, and were enriched in putative venom components (Dennis *et al.* 2017).

**Figure 1.**
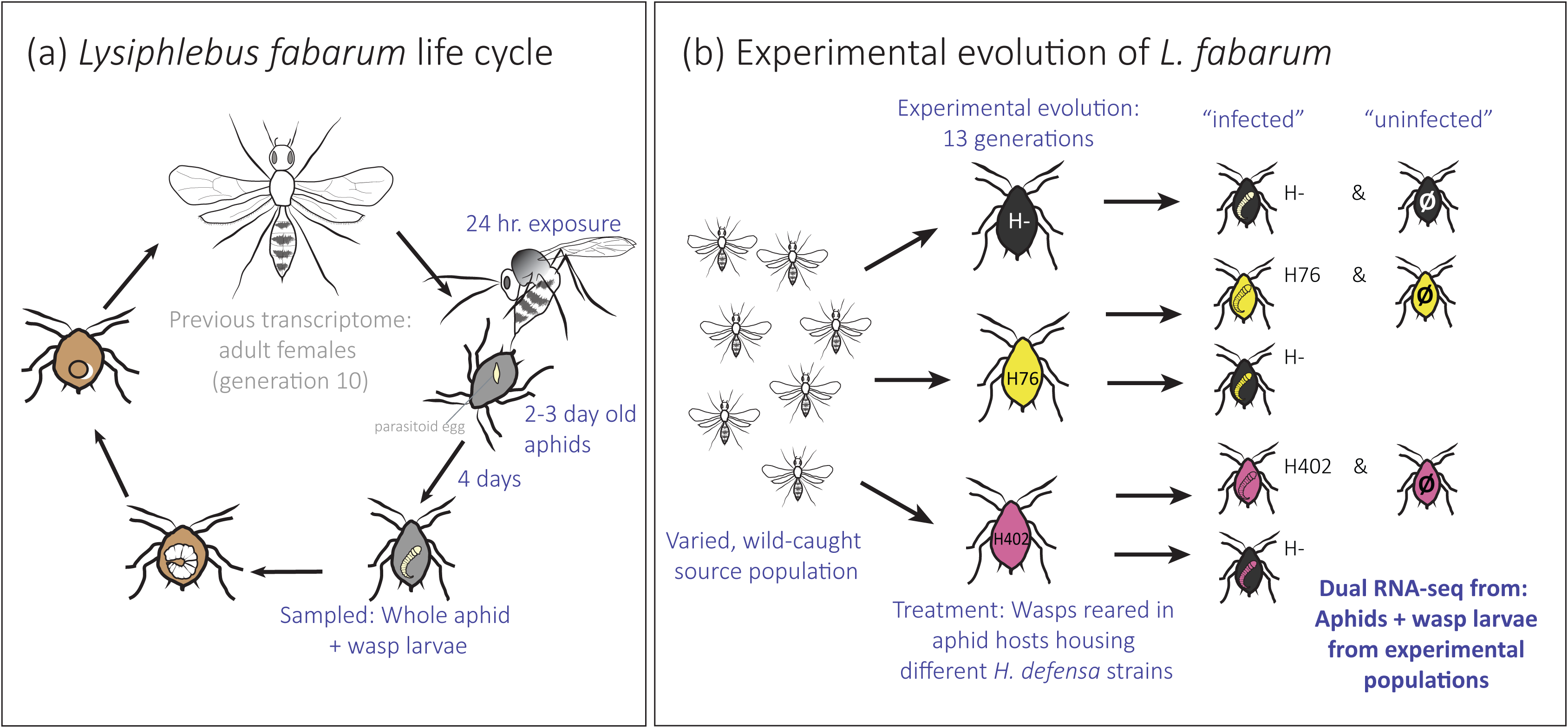
(a) The life cycle of *L. fabarum* and sampling design for RNA-seq (b) Summary of experimental evolution of *L. fabarum* reared in different aphid hosts. Each of the three treatments was replicate in four independent populations. All lineages were sampled from the subline that hosted them during experimental evolution and H- aphids. Aphids marked Ø were deemed free of parasitoid infection prior to sequencing.

However, *H. defensa*-conferred resistance does not target the adult wasps, it targets the eggs or developing larvae inside the aphids’ bodies. To better study this, the analysis of gene expression should ideally capture both aphid host and parasitoid together. Here, we have employed dual RNA-seq (Westermann *et al.* 2016; Westermann *et al.* 2012; Wolf *et al.* 2018) to examine gene expression in 4-5 day old parasitoid larvae within still living aphids. We have identified gene expression changes in both aphids and their infecting parasitoids, observed dramatic shifts in expression associated with infection success, and identified suites of parasitoid genes associated with both venom and detoxification that are differentially expressed among lineages with different infective abilities.

## 2. Materials and Methods

### 2.1 Insects

The black bean aphid (*Aphis fabae*) is a widely distributed agricultural pest found across Europe (Blackman & Eastop 2006). The three aphid sublines used in this study were established from a single aphid clone (Vorburger *et al.* 2009) and have been maintained in the lab for >100 generations. We utilized three sublines: two that are protected by the endosymbiont *H. defensa* (A06-407^H402^ and A06-407^H76^, hereafter H402 and H76) and one unprotected subline (A06-407, hereafter H-). The protective *H. defensa* were chosen for their distinct genotypes in two housekeeping genes and the varying levels of protection they provide against multiple lines of the parasitoid *L. fabarum* (Cayetano *et al.* 2015). The APSE possessed by these *H. defensa* strains encode different toxins: H76 encodes a YD-repeat that provides high protection against parasitoids, while H402 encodes a cytolethal distending toxin (CytD) that provides medium protection to aphids (Brandt *et al.* 2017; Cayetano *et al.* 2015; Dennis *et al.* 2017).

The parasitoid wasp *L. fabarum* (Hymenoptera: Braconidae: Aphidiinae) is the most important parasitoid of *A. fabae* in the field (Rothacher *et al.* 2016) and one of the most abundant parasitoids in Europe (Starý 2006). Females inject eggs into the aphid hemocoel, and development proceeds until a single wasp pupates and emerges as a free-living adult. During this development the aphid is eventually killed and wasp development completes in the “mummy” exoskeleton that remains.

The *L. fabarum* lineages studied here were established by experimental selection, and were founded from a diverse, wild-caught population, described in depth in Dennis *et al.* (2017). Briefly, wasps were collected from six sexually reproducing populations across Switzerland, each maintained separately in the lab for 24-30 generations and then combined in a large pool for two generations. This common, genetically variable stock population was used to establish founding populations of 50-60 individuals for experimental evolution, each housed on one of the three aphid-hosts described above (hereafter: treatments H76, H402 and H-). Each treatment was independently replicated four times. All populations were maintained throughout the experiment at 22°C and with a 16-hr photoperiod. Each replicate was reared on three broad bean plants colonized by the respective aphid sublines; aphids were replaced from a clonally reproducing, un-parasitized stock each generation, thus removing their ability to adapt to parasitoids.

### 2.2 Sampling

To simultaneously examine expression in growing parasitoid larvae and their aphid hosts, we exposed 3-4 day old aphids to wasps from each of the 12 experimentally evolved populations and collected them four days later (i.e. aphids were 7-8 day old at the time of sampling, Figure 1). We used parasitoids following 14 generations of experimental evolution. At this time-point the H76 and H402 treatments were significantly better at infecting the aphids on which they had been reared, but had not improved their abilities to infect the other two aphid sublines (Dennis *et al.* 2017). Whole aphids were frozen directly at −80 °C at 14:00hr; all aphids were alive when sampled. To establish which aphids held successfully developing wasp larvae, we performed simultaneous RNA-DNA extractions with Trizol reagent (Invitrogen, Carlsbad, CA, cat #15596) according to the manufacturer’s instructions but for an overnight incubation in Trizol. Freshly extracted RNA was stored at −80 °C, while DNA from the same sample was screened for parasitoid presence with two species-specific microsatellite markers developed for *L. fabarum* (Lys2 and Lys5; Sandrock *et al.* 2007), and one species-specific aphid microsatellite primer (AF-85, Coeur d’acier *et al.* 2004). Samples were screened on agarose gels; samples testing positive for *Aphis* and negative for both *Lysiphlebus* primers were deemed un-successful parasitoid infections. Only samples with positive bands for all three primer pairs were deemed successful.

We aimed to examine evolved differences among lineages separately from the plastic, short-term response to the presence of *H. defensa*. As shown in Figure 1, we achieved this by rearing parasitoids from all 12 experimentally evolved populations (four replicates per treatment) under their experimental conditions (e.g. parasitoids from treatment H402 reared in aphid subline H402), and by rearing parasitoids experimentally evolved in the presence of *H. defensa* (i.e. treatments H402 and H76) in *H. defensa*-free aphids (H-) as well. We aimed to sequence three biological replicates (i.e. separate individuals) from each population, but low parasitism rates did not make this possible in all instances (Supplementary Table 1). We also chose three “uninfected” aphids from each treatment, although in all cases parasitoids could have attempted to infect these individuals. In total, 385 aphids were screened for wasp presence and 51 individuals were chosen for RNA-seq, including nine apparently uninfected aphids (Figure 1, Supplementary Table 1).

RNA from samples chosen for transcriptomic sequencing was further purified using the RNeasy mini kit (Qiagen, Venlo, Netherlands, cat # 74104), verified on an Agilent 2100 Bioanalyzer (Agilent Technologies, Santa Clara, CA, USA), and quantified with a Qubit 2.0 Fluorometer (Life Technologies, Carlsbad, CA, USA). Stranded cDNA libraries were constructed using half reactions of the Illumina TruSeq Stranded mRNA kit (#RS-122-2101) according to the manufacturer’s instructions, with an input of approximately 500ng total RNA. Libraries were quantified by qPCR and randomly assigned to four lanes for 100bp, single-end sequencing on an Illumina HiSeq2500 (v3 chemistry) at the Functional Genomics Center Zürich.

### 2.3 *de novo* transcriptome assembly

Raw Illumina reads were quality filtered to remove all sequences with an “N” in Prinseq (Schmieder & Edwards 2011), followed by primer trimming and quality filtering (q<20, 5bp sliding window) with Trimmomatic (Bolger *et al.* 2014). Read quality was summarized with FastQC (Andrews *et al.* 2010).

For read mapping, we constructed two separate references for *L. fabarum* and *A. fabae*. Transcripts belonging to *A. fabae* were *de novo* assembled from the nine aphids collected here that were deemed wasp-free by PCR screening (above) and four additional libraries constructed from *H. defensa*-free aphids from the same aphid clone, also 7-8 days old, that had never been exposed to parasitoids (Nation Center for Biotechnology Information (NCBI) Sequence Read Archive (SRA): SAMN10606880, SAMN10606885, SAMN10606890, SAMN10606895, SAMN10606900, SAMN10606905, SAMN10606910, SAMN10606915, Käch unpublished). These latter libraries were prepared and quality filtered using the same methods as in this study, except that instead of poly-A enrichment they were depleted of ribosomal RNA using half reactions of the riboZero Epidemiology kit (Illumina, MRZE706) and were paired-end sequenced.

The *L. fabarum* reference transcriptome was assembled in two steps. First, we used the transcriptome previously constructed from >370 adult females, in 59 separate Illumina libraries, from these same experimental evolution lines at generation 11 (raw data under NCBI SRA PRJNA290156, Dennis *et al.* 2017). Second, to incorporate transcripts not present in the previous transcriptome, we mapped the 51 new libraries generated here to this reference, extracted reads that did not map, and constructed novel *L. fabarum* transcripts using the *L. fabarum* draft genome (available from: bipaa.genouest.org/is/parwaspdb; Dennis *et al.* in prep) as a guide in Trinity v2.4.0 (Grabherr *et al.* 2011).

To ensure that the two transcriptomes contained only their intended species, we used *blastn* (Camacho *et al.* 2009) to assign transcripts to five sources that were most likely present in this data: (1) the *L. fabarum* draft genome assembly (Dennis *et al.* in prep), (2) *A. fabae* scaffolds from three different genome assemblies from other *Aphis* species [*A. gossypii* scaffolds(bipaa.genouest.org/is/parwaspdb) and *A. glycines* v1 (Wenger *et al.* 2017) and *A. glycines* Biotype1 (bipaa.genouest.org/is/parwaspdb)], (3) the defensive symbiont *H. defensa* isolated from *Acyrthosiphon pisum* (Accession CP001278, Degnan *et al.* 2009), (4) the obligate symbiont *Buchnera aphidicola* strain BAg, isolated from *Aphis glycines* (Genbank NZ_CP009253, NZ_CP009254, and NZ_CP009255, Cassone *et al.* 2015), and (5) the human genome (hg38, Accession: GCA_000001405.26). Transcripts were assigned to a species by megablast (e-value > 1e-5), and sequences matching to more than one taxon were retained only if the bitscore gain of the best matching taxon was greater than 100; otherwise the transcript was discarded. All transcripts that did not match to *L. fabarum* or *A. fabae* with these criteria were discarded. Remaining transcripts were clustered with CD-HIT-EST (Li & Godzik 2006) to remove redundancy (98% threshold), performed separately within each species. To ensure that we did not analyze reads that mapped to both species, the final *L. fabarum* and *A. fabae* transcriptomes were joined into a single reference for mapping.

### 2.4 Differential expression (DE) analysis

Read counts were generated by mapping the individual libraries to the combined *A. fabae* and *L. fabarum* transcriptome using RSEM (Li & Dewey 2011), and bowtie2 (Langmead & Salzberg 2012), employed via the “align_and_estimate_abundance.pl” script packaged with Trinity. From this, gene-level count estimates were imported in R (R-Core-Team 2012) to DESeq2 (Love *et al.* 2014) using tximport (Soneson *et al.* 2015), which corrects for length differences among isoforms and reduces biases that could arise when isoforms with different lengths are expressed in different treatments. The resulting counts were analyzed for differential expression based on a negative binomial distribution of read-counts using DESeq2. Importantly, this pipeline provides DESeq2 with gene-level counts, an approach that has previously been shown to have advantages over transcript-level analyses (Soneson *et al.* 2015). In all analyses, the models included individual replicate lines nested within treatment, and analyzed aphid and wasp reads separately. Read counts were filtered prior to the analysis to retain genes with at least ten reads in a third of the included samples. We only included individuals with at least 1 million raw reads; because of unequal expression between host and parasitoid, different libraries were discarded for wasps and aphids (Supplemental Table 1).

We analyzed data at several levels to examine both overall responses to infection and lineage-specific responses to parasitism. For clarity, we have labeled these levels of analysis: Analysis 1, 2, and 3. In Analysis 1, we examined the overall impact of infection in both parasitoids and aphids across the entire dataset of 51 libraries, and based our designation of infection success on our PCR-based wasp detection (See results Figure 2). This revealed two distinct clusters of samples in the analyses of both aphids and wasps, but the “uninfected” group included several individuals that we expected to be “infected”. Because our PCR-based detection was only an estimate of wasp success, we concluded that this large expression difference was a better determinant of wasp success than PCR, which could have amplified trace DNA from dead or dying wasps. This means that seven additional aphids were deemed “uninfected” and were excluded from all further analyses, as these only examined ongoing infections.

**Figure 2.**
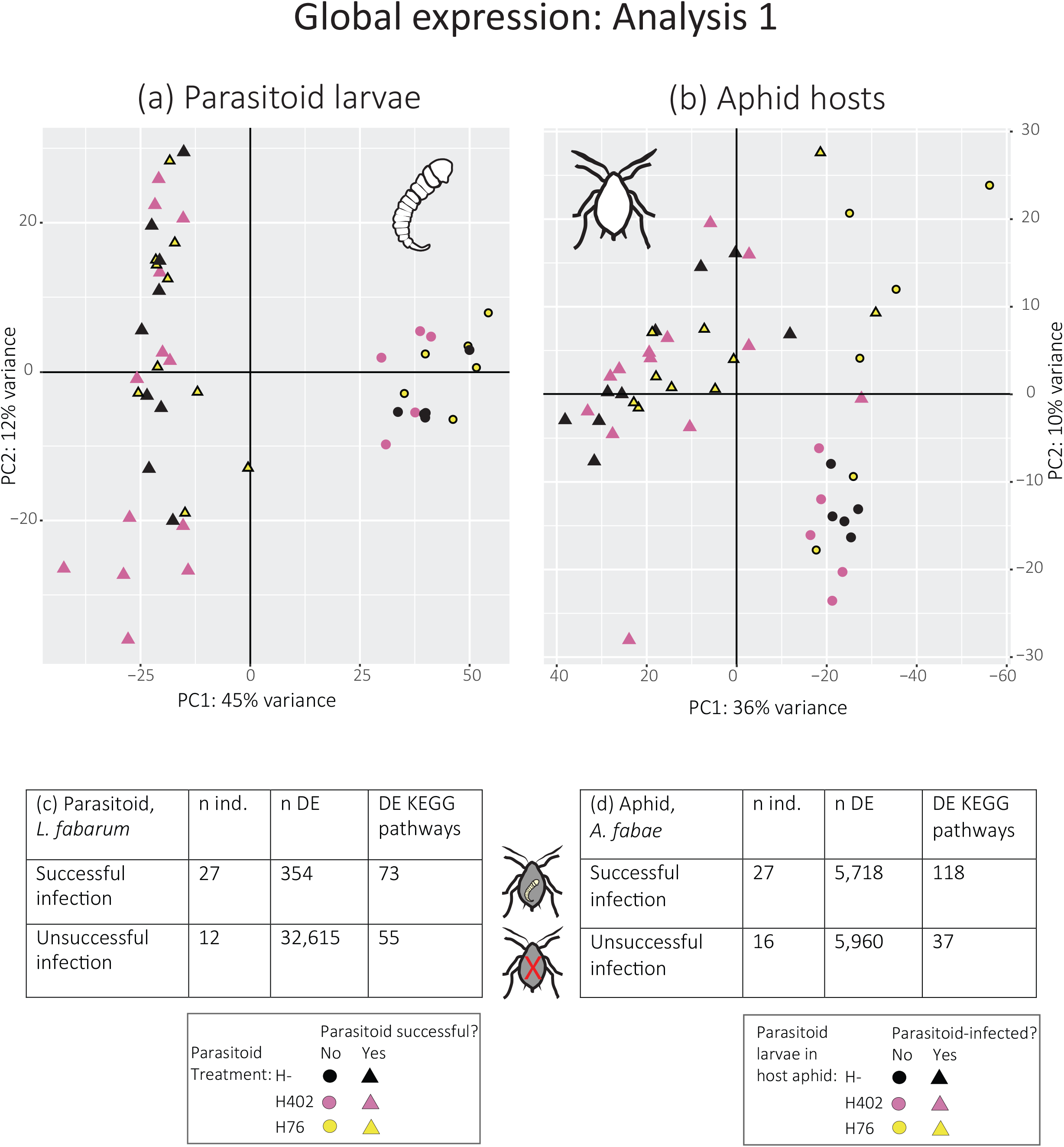
Expression patterns across all samples: (a) Global expression profile in parasitoid larvae, including successfully infecting (triangles) and failed (circles) individuals. Symbols are colored according to experimental evolution treatment, (b) Global expression profiles for host aphids, including infected (triangles) and uninfected (circles) individuals, (c) summary of analysis of differential expression between successful vs. failed parasitoid larvae, and (d) infected vs. uninfected aphids. Abbreviations. number of individuals sampled (n ind), number of differentially expressed genes (n DE), and differentially expressed KEGG pathways (DE KEGG pathways).

In Analysis 2, we compared expression among only the endosymbiont-free (H-) aphids and the larvae they contained (Figures 3 and 4). In aphids, this means we compared identical, *H. defensa*-free clones, infected with parasitoid larvae from the three different treatments. This allows us to detect any response of the aphids themselves, and not their endosymbionts, to the potential differences in infective strategy among parasitoids. In parallel, we compared expression among the parasitoid larvae. By rearing the three parasitoid treatments for one generation in endosymbiont-free aphids, we aim to avoid short-term responses to the endosymbionts present in defended aphids. Thus, we interpret expression differences in Analysis 2 as largely genetically-based, evolved differences among the experimental parasitoid populations. We also re-analyzed previously collected adult female expression in samples collected from the same lineages and experimental design (i.e. H- aphids, Figure 4), collected three generations prior (generation 11, Dennis *et al.* 2017). By re-analyzing these individuals, we sought to examine overlaps in expression between adult and larval wasps. In all three comparisons (aphids, larval wasps, and adult wasps), we modeled expression in DESeq among treatments and examined the pairwise expression differences between the H402, H76, and H- treatments.

**Figure 3.**
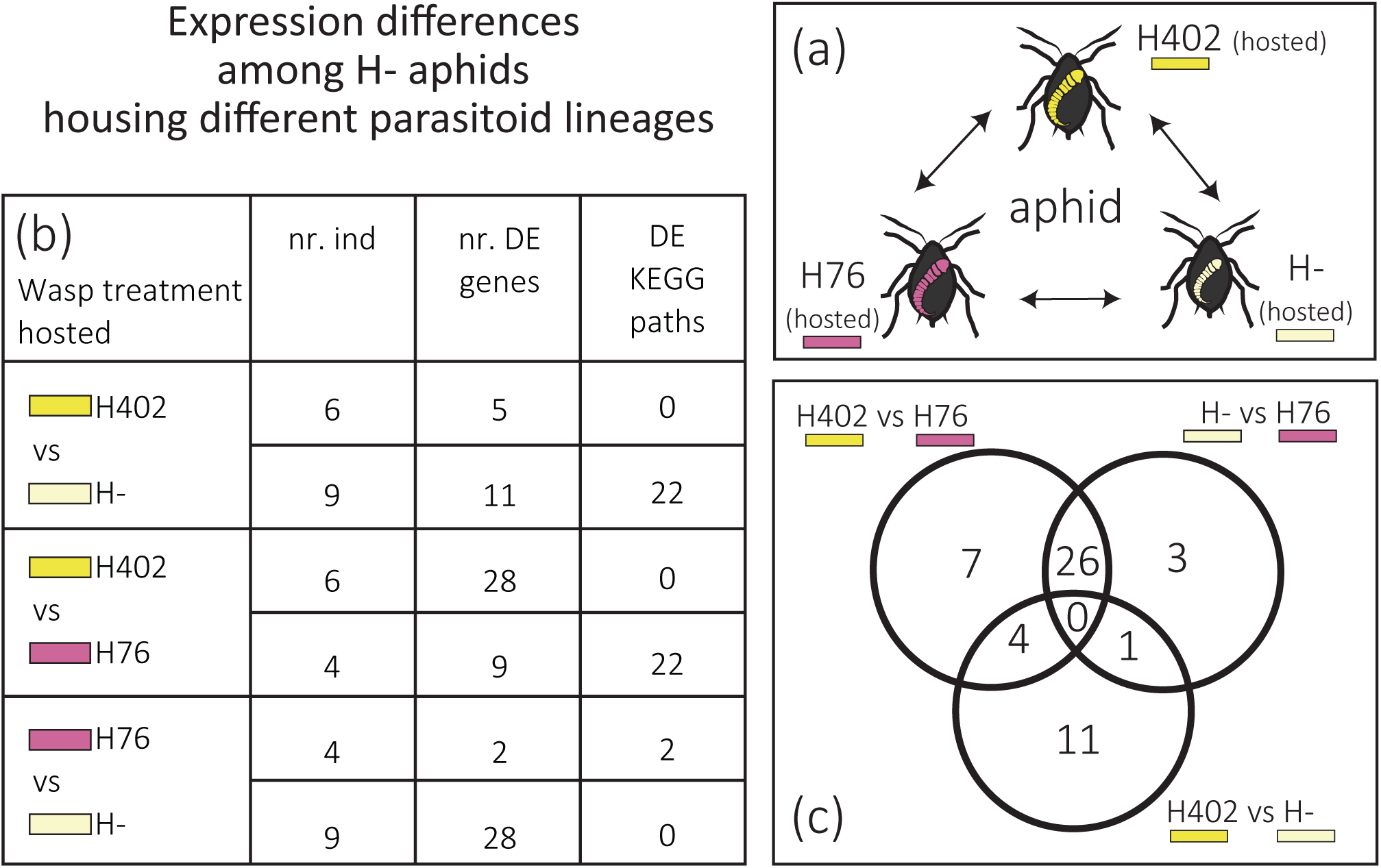
(a) Analysis of differential expression in H- aphids possessing parasitoid larvae from the three experimental evolution treatments, (b) summary of pairwise comparison from model of differential expression, and (c) the full model of differential expression was summarized in three pairwise comparisons and there was some overlap among these comparisons. Abbreviations: Number of individuals sampled (nr. ind), number of differentially expressed genes (nr. DE), and differentially expressed KEGG pathways (DE KEGG paths).

**Figure 4.**
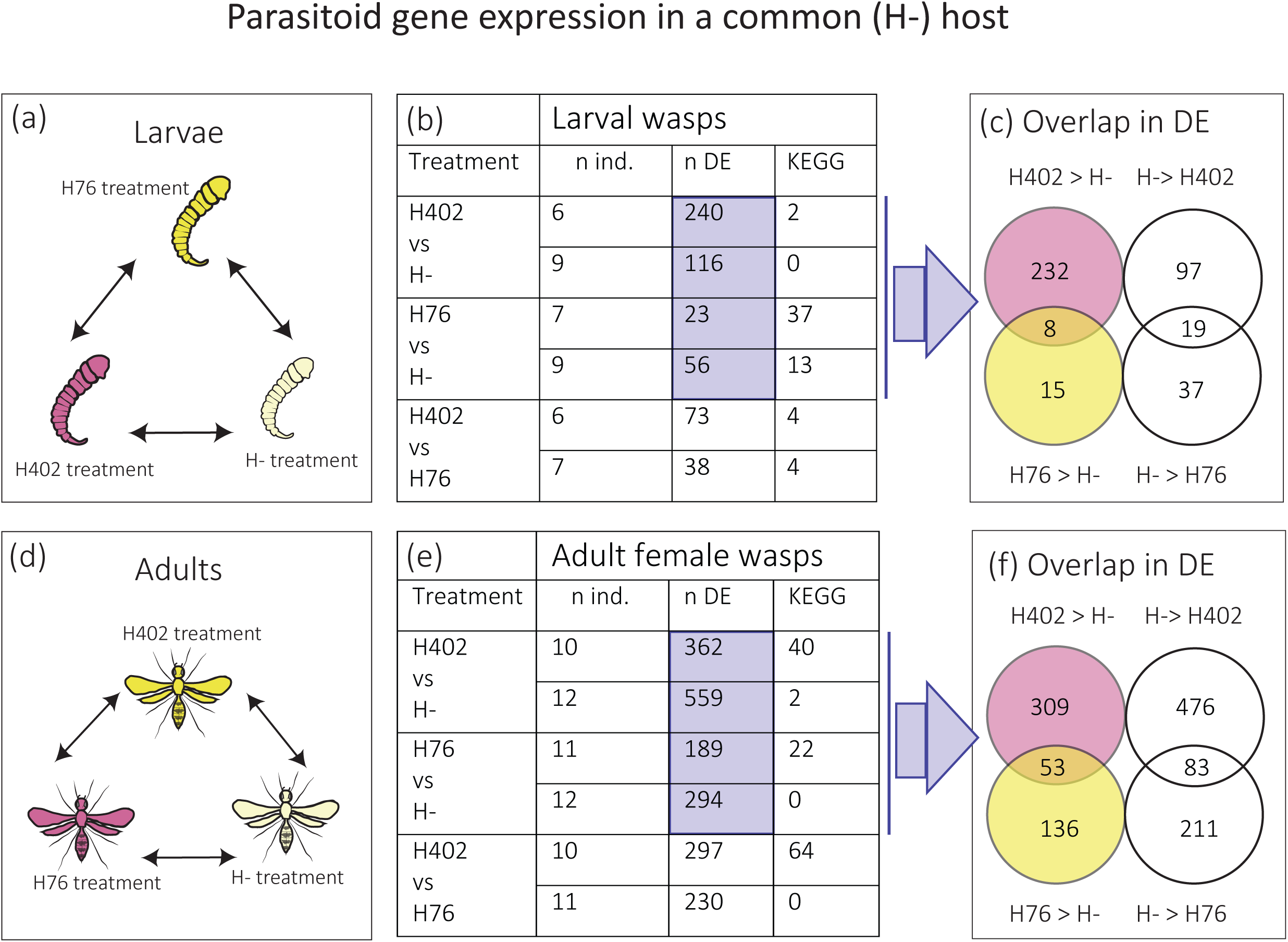
Analysis of differential expression in (a) larval and (d) adult female wasps following experimental evolution. Summary of differential expression and KEGG results for (b) larvae and (e) adults were analyzed in two separate models, each summarized by three pairwise comparisons. Venn diagrams (c, f) show the overlap in the differentially expressed genes from treatments reared in the presence of *H. defensa* (H402 or H76); these overlapping genes represent genes expressed to cope with *H. defensa* in general. Note: in adult females, six transcripts were DE in opposite directions in this Venn diagram; this is not shown here and has been previously reported in Dennis et al (2017). Abbreviations: Number of individuals sampled (nr. ind), number of differentially expressed genes (nr. DE), and differentially expressed KEGG pathways (DE KEGG paths).

Lastly, Analysis 3 examined plastic responses to *H. defensa* separately within lineage by comparing the H76 or the H402 lineages reared in either H- or their respective H+ (H76 or H402) hosts. We did not compare aphid expression in this subset of the data because we expect that differential expression will result mostly from the presence and absence of *H. defensa*, and this was the focus of a separate investigation into the impact of many different *H. defensa* strains on aphid expression (Käch unpublished).

### 2.5 KEGG pathway enrichment

To examine pathway-level differences in gene expression, we used the Kyoto Encylopedia of Genes and Genomes (KEGG: Kanehisa & Goto 2000) within both Analyses 1 and 2, for aphids and wasps. For this, we used the KEGG Automatic Annotation Server (Moriya *et al.* 2007) to assign transcripts to pathways, and analyzed expression among these using the Generally Applicable Gene-set Enrichment GAGE (Luo *et al.* 2009) and Pathview (Luo & Brouwer 2013) packages in R. We broadly examined gene expression differences by combining read counts across replicates within treatment. Raw reads were log2 transformed and normalized to account for unequal sizes among libraries, and we examined significant pathways with an adjusted p-value (FDR) < 0.1.

## 3. Results

### 3.1 Dual RNA-seq reference transcriptome and read mapping

In total, we generated 1.59×10^9^ single-end reads (NCBI SRA Accessions: SAMN10024115-SAMN10024165), >99% of which remained after quality filtering. The *A. fabae* reference transcriptome was generated from 1.99×10^8^ of our single-end reads (our nine “uninfected” aphid samples) and 1.34×10^8^ paired-end reads from the same aphid clone, never exposed to wasps. After removing ribosomal sequences that should have been eliminated in the lab (28S and 18S), this generated 250,745 transcripts for *A. fabae*; we speculate that this transcriptome is quite large because it was constructed from whole aphids, which house many different tissues, including reproductive tissues containing two subsequent generations of offspring. For *L. fabarum*, we generated 106,516 transcripts. Thus, the joint transcriptome to which we mapped our 51 quality-filtered libraries contained 357,261 transcripts (Supplementary Table 1).

Initial examination of the data revealed that many reads did not map to either species. Further investigation revealed that the major portion of these lost reads mapped to just six transcripts that did not *blast*-match to either focal species (Supplemental Table 1, 2). Based on their best *blast* match, we believe five of these belong to a novel group of viruses that have thus far been dubbed ‘dark matter’ viruses (Supplemental Table 2). These abundant viruses are not easily classified because they are highly divergent and rapidly evolving, but these sequences are similar to several segments recently identified from *Drosophila* (Krishnamurthy & Wang 2017; Obbard 2018; Webster *et al.* 2015). The sixth sequence matched to a positive stranded RNA virus [ssRNA(+)] virus (Supplemental Table 2). Viral load varied among samples (Supplemental Table 1), and totaled >480 million reads. Viral reads of all types were not included in any analyses of differential expression. Excluding these viral sequences and other contaminants, the tximport-weighted mapped reads totaled 334 million for *L. fabarum* and 444 million for *A. fabae.* One library was excluded from all analyses because the PCA of all expressed genes showed it to be a strong outlier (Supplemental Table 1).

### 3.2 Analysis 1: Comparative gene expression across all samples

#### 3.2.1 Response to infection in aphids

Overall expression patterns among aphids (Analysis 1) separated them into two clear groups of infected and uninfected; this included seven individuals that were newly designated uninfected based on this data (Figure 2, Supplemental Data 1). Over 10,000 genes were differentially expressed and these were nearly evenly split into genes that were more highly express in infected and uninfected aphids (Figure 2). Within these, >100 KEGG pathways were differentially expressed, many of which suggest that infected aphids increase metabolic processes. There were 28 metabolic pathways upregulated in infected individuals, including upregulation of Carbon, Glycine, Sucrose, Methane and Pentose metabolism (Supplemental Data 2). Pathways implicated in immunity were also upregulated in infected aphids, including secondary metabolite (ko01110) and antibiotic (ko01130) biosynthesis, two components of the toll pathway (Toll-like receptor signaling pathway, ko04620 and Toll and Imd signaling pathway, ko04624), T cell receptor signaling pathway (ko0460), and the NOD-like receptor signaling pathway (ko04621). KEGG pathways that were more highly expressed in uninfected aphids do not include any metabolic paths, but do cover cell cycle and pathways associated with growth (e.g. DNA replication, Hippo signaling).

#### 3.2.2 Differential expression in infecting parasitoid wasps

Even when parasitoid infection was deemed unsuccessful, aphids yielded some parasitoid RNA, presumably from dead or dying larvae. This allowed a comparison also for parasitoid wasps. The separation in overall expression patterns between successful and unsuccessful infections was even more distinct than in aphids, and the two groups contained the same individuals as in aphids (Figure 2). Unexpectedly, there are many more genes that are statistically identified as upregulated in unsuccessful (32,615) than in successful larvae (354, Figure 2, Supplemental Data 3). However, examination of this data shows that this is almost entirely a product of large spikes in expression of some genes in just one or few individuals (Supplemental data 3). This large upswing in expression is also not well organized, as fewer KEGG pathways were upregulated in dead and dying larvae (55 pathways) than in successful larvae (73 pathways, Supplemental Data 4).

Unsurprisingly, pathways upregulated in successfully developing wasps include basic biological functions such as Oxidative Phosphorylation (ko00190), the TCA cycle (ko00020), and Cell cycle (ko04110). Successful wasps also upregulated genes associated with metabolism (16 in total, including: Amino sugar and nucleotide sugar metabolism, Fatty acid metabolism, Fructose and mannose metabolism). In contrast, only one metabolic pathway (Tyrosine, ko00350) was upregulated in unsuccessful wasps. Among successfully infecting wasps, one of the most highly supported upregulated pathways is proteins associated with the Ribosome (ko03010). This KEGG term encompasses a number of ribosomal accessory proteins, and many of these were also individually differentially expressed; at the gene level there are at least 130 differentially expressed ribosomal genes, and these comprise nearly 20% of all genes upregulated in successful parasitoids.

### 3.3 Analysis 2: Evolved expression differences within successful infections

#### 3.3.1 Gene expression in H- aphids hosting different parasitoids

Analysis 2 compared expression among symbiont-free aphids while housing the three parasitoid treatments. Here, the overall analysis modelled expression among the (genetically identical) aphids containing the three wasp treatments, and yielded results in three pairwise comparisons. There were relatively few (2-28 genes, Figure 3) significantly differentially expressed genes in this comparison (Figure 3, Supplementary Data 5). Unfortunately, of the few results that we did find not all could be identified by our *blast* annotation (Supplementary Data 5). Among the genes that could be identified, expression was more similar between the aphids hosting H- and H76 parasitoid treatments (26 shared results, Supplemental Data 5), than either group relative to the H402 treatment. Several putative transcription modifiers and gene regulators are more highly expressed in aphids housing the H-parasitoid lineage, relative to H76. Aphids housing parasitoid larvae from the H76 lineage show higher expression of a chitinase. Aphids housing H402 larvae express higher levels of a ubiquitinase, a phosphatase, and a putative sphingomyelin synthase, which may regulate endoplasmic reticulum production.

KEGG pathway analysis (Supplementary Data 6) offers further support of activity relating to protein processing; aphids housing parasitoids from the H76 treatment have higher expression of just two KEGG pathways, matching to Protein export (ko03060) and Protein processing in the endoplasmic reticulum (ko04141). Pathways that are more highly expressed in aphids infected by both treatments H- and H76 (each relative to H402-housing aphids) include Biosynthesis of antibiotics (ko01130), Biosynthesis of secondary metabolites (ko01110), and a suite of metabolic pathways (Supplementary Data 6).

#### 3.3.2 Expression differences among parasitoid lineages in a common host

Analysis 2 also examined expression among parasitoid larvae from the three evolution treatments when reared in a common (H-) host environment (Figure 4, Supplemental Data 7), and generated three pairwise comparisons to mirror the aphid analysis. Differential expression among parasitoid treatments was largely specific to that treatment, as was previously observed in adult females from these lineages (Dennis *et al.* 2017). Only 28 transcripts (nine up and 19 down, Figure 4) were differentially regulated in both H76 and H402 relative to the H- treatment. Only four of the upregulated genes are annotated, and these appear to be involved in regulation of expression (e.g. two are Kreuppel-like factors, likely transcription repressors, Supplemental data 7). Of the shared genes upregulated in the H- treatment (higher than both H402 and H76 treatments), only nine are annotated; three of these are heat-shock proteins (HSP 68/70) and one is a ribosomal protein (L19, Supplemental data 7). These likely represent the general mechanisms by which parasitoids have evolved to cope with the presence of *H. defensa*.

A remaining 375 differentially expressed larval genes are specific to just one lineage (Figure 4, Supplemental data 7). Among these, we observe genes with several key functions, and of particular interest are those that are detoxification/stress-associated genes or venom/toxicity genes. Genes likely to help detoxify or cope with stress include seven additional heat-shock proteins that are more highly expressed in the H- treatment (Hsp 68, Hsp 70, lethal(2)essential/Hsp20) and three different genes (Hsp 70 and lethal(2)essential/Hsp20) that are upregulated in the H402 treatment. Numerous ribosomal proteins (L19, L21, L4, L35, and S18) are differentially expressed between the H- and H402 treatments. As previously observed in adult females we see differential expression of putative venom or toxic compounds (Dennis *et al.* 2017). These include five metallo-endopeptidases and neprilysins (all more highly expressed in H402 treatment), and three leucine-rich-repeat proteins (more highly expressed in the H- treatment). However, there are fewer candidate venom toxins than in adult females; this makes sense given that larvae are not expected to produce venom, and given that some of these samples are likely male. A number of transcriptional regulators are also differentially expressed in each lineage, as well as histones (upregulated only the H402 treatment). While we cannot yet determine the genes with which these interact, they support the idea that regulatory differences help drive expression differences between parasitoid treatments.

KEGG pathways that are more highly expressed in the H76 treatment relative to H- include Ribosome (ko03010), Biosynthesis of antibiotics (ko01130), and a number of pathways suggesting higher metabolism of wasps evolved in the presence of *H. defensa* strain H76 (e.g. Oxidative phosphorylation, Carbon metabolism, TCA cycle, Supplementary Data 8). The Ribosome, Biosynthesis of Antibiotics, and Oxidative phosphorylation pathways are also upregulated in H76 relative to H402. In treatment 402, there are no pathways that are more highly expressed: only two are downregulated relative to the H- treatment: Proteasome (ko03050) and Tyrosine metabolism (ko003500). In relation to the H76 treatment, H402 upregulated pathways suggesting growth (Hippo signaling, Axon guidance and DNA replication).

We compared these results to samples taken from the same experimental design in adult females from the same lineages, newly analyzed with the extended reference transcriptome (Figure 4, Supplemental Data 9). Previous analyses already addressed differences among the three treatments in a common environment (Also called Analysis 2: Dennis *et al.* 2017), so we will only discuss the analysis of this data in comparison to the larvae (Supplementary data 10). To examine expression differences in both larvae and adults, we looked at all genes that were differentially expressed in more than one comparison (remembering that there are six pairwise comparisons across the two analyses; Figure 5). There were 42 genes that were differentially expressed in both of the life stages. While not all genes were identified, there were several interesting groups (Figure 5). Heat shock proteins (Hsp68 and Hsp70, three genes) were upregulated in the H- treatment relative to both H402 and H76; three different genes identified as Hsp70 were also upregulated in the H402 treatment. Three different transcripts identified as NEDD4-binding protein-2 are upregulated in only the H402 treatment. Two ribosomal proteins (S18 and L35) are more highly expressed in the H402 and H- treatments. A putative venom component (F-box leucine-rich-repeat protein) was more highly expressed in the H- and H402 treatments.

**Figure 5.**
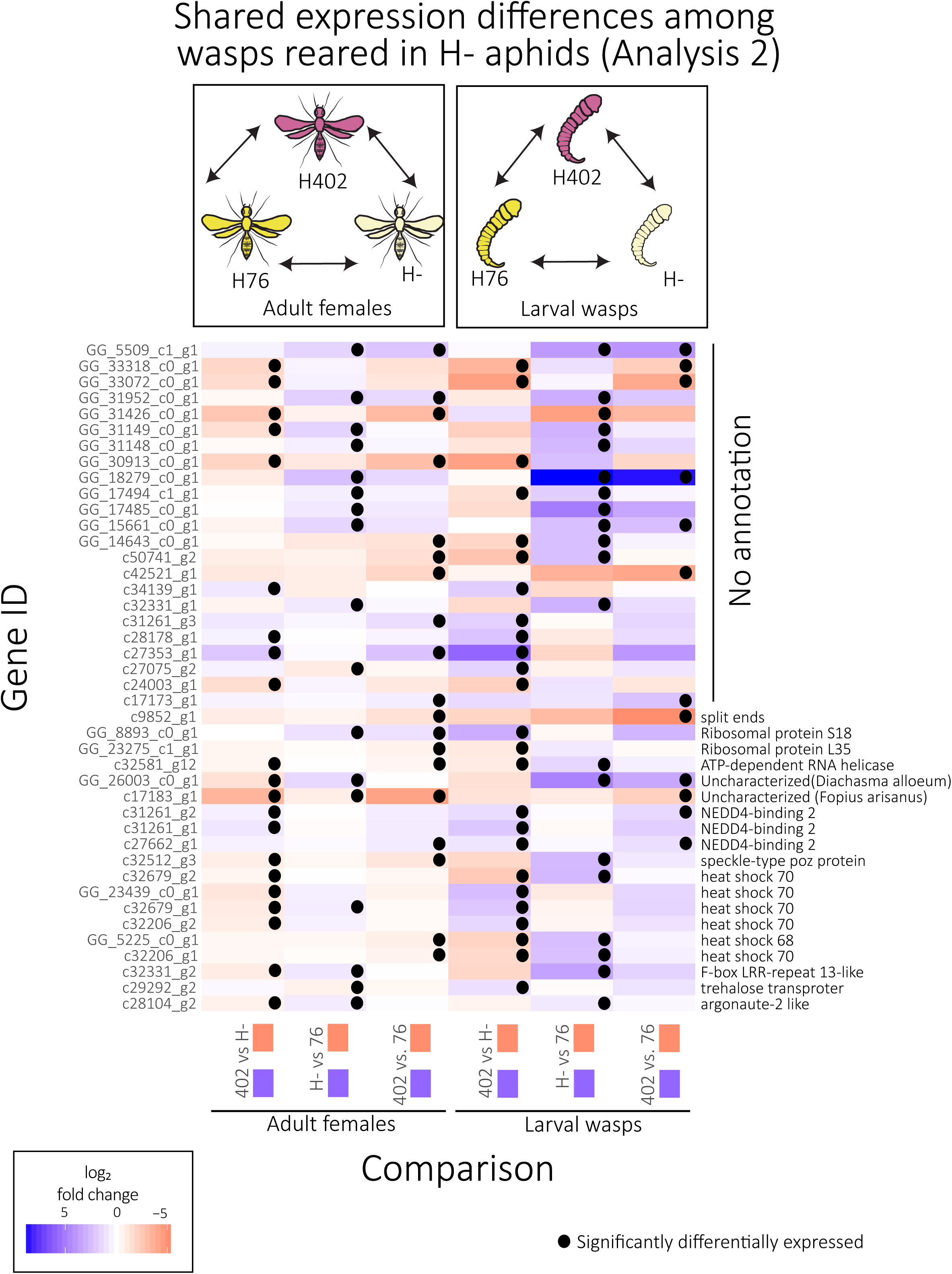
Heatmap depicting log_2_(fold change) of genes that were differentially expressed in parasitoids at two life stages. In both datasets, the analysis included only parasitoids reared for one generation in a common aphid environment (without *H. defensa*). Black dots indicate significant expression changes. Direction of change is matched to the treatment by the colored box below the x-axis.

### 3.4 Analysis 3: Plasticity in parasitoid gene expression

Within the H76 and H402 treatments, we used separate comparisons of expression between larvae collected from aphids that were symbiont free (H-) or not (H76 or H402, respectively). We did not make a parallel comparison within the host aphids, because their biology is known to be largely impacted by the presence or absence of protective endosymbionts, and this has been addressed in other specific studies (Cayetano *et al.* 2015; Käch unpublished; Vorburger & Gouskov 2011).

#### 3.4.1 Plasticity in parasitoid treatment H402

With treatment H402, we identified 20 differentially expressed genes between individuals reared on H- or H402 aphids (Table 1). None of the six genes that were more highly expressed in the presence of *H. defensa* were *blast* annotated (Supplementary Data 11). Of interest among the genes upregulated in H- hosts (only four were annotated) were one cuticular protein and a translation initiation factor.

**Table 1:** Plastic response in parasitoid gene expression. Comparison of either treatment H402 or treatment H76 larvae, hosted on by aphids with or without their respective *H. defensa*.

| Analysis 3: Plastic responses |  |  |  |
| --- | --- | --- | --- |
| Parasitoid lineage | Aphid host | n individuals | n DE |
| Treatment H402 | H402 vs. | 5 | 6 |
|  | H- | 6 | 14 |
| Treatment H76 | H76 vs. | 3 | 24 |
|  | H- | 7 | 9 |

#### 3.4.2 Plasticity in parasitoid treatment H76

Unfortunately, several of the libraries from the H76 treatment had low read counts (Supplemental Table 1); nonetheless we present these results to show that there are likely small plastic responses to the toxic environment created by *H. defensa*. We identified 33 differentially regulated genes in the H76 treament (Table 1). These include several candidates identified in previous comparisons, that appear to be further upregulated when these lineages are in the presence of *H. defensa*, including ribosomal protein L4, a small heat shock protein (lethal(2)essential for life), and two leucine-rich-repeat proteins (Supplemental Data 12).

## 4. Discussion

### 4.1 Gene expression globally shifts in response to parasitoid infection

Parasitoid infection had a clear impact on gene expression in host aphids (Analysis 1, Figure 2). The most striking pattern in aphids housing growing wasp larvae was an increase in metabolic processes, illustrating energy demands and a redirection of metabolic pathways that is unsurprising given the changes this invokes in the aphid (Beckage & Gelman 2003; Starý 1970). Key immune pathways were also upregulated in infected aphids, including the Toll pathway. The KEGG pathway associated with biosynthesis of antibiotics is also upregulated in infected aphids. Although it would require further investigation, we raise the possibility that infected aphids could be losing their *H. defensa*, in a process analogous to destabilization and bleaching in corals (Coles & Brown 2003), or that these genes have a different function in this system.

More surprising was the upregulation of numerous genes in failing parasitoid larvae, but we are quite confident that this is largely a statistical artifact resulting from disorganized spikes in gene expression of dying larvae. These could be symptoms of decline or even death; organized gene expression has been documented up to three days after death, and has included genes associated with development, stress, and transport (Pozhitkov *et al.* 2017), all of which are also upregulated here. Importantly, this supports other work suggesting that even when parasitoids do not successfully infect, they can still have long term impacts on aphid survival and reproduction (Vorburger *et al.* 2008) and leave detectable amounts of mRNA, although the negative impact could be compensated in the following generation (Kaiser & Heimpel 2016).

Despite the differences among parasitoid treatments, the most striking pattern of differential expression across all successful larvae is that 70 out of the 354 upregulated genes are ribosomal proteins (Supplemental Data 3). Ribosomal proteins surround the primary ribosomal components and impact translational choice and efficiency (Brodersen & Nissen 2005; de la Cruz *et al.* 2015). Upregulation of many of these same ribosomal proteins in eukaryotes has been associated with tumor suppression, immune signaling, development and disease response (Zhou *et al.* 2015). A growing number of studies have identified upregulation of ribosomal genes in association with a variety of environmental stresses, including dehydration (in rice: Moin *et al.* 2017), predator-induced stress (Hales *et al.* 2017), and fluoride stress (Tang *et al.* 2018). Thus, upregulation of ribosomal proteins may be a widespread and important response to stress, and the evolution of their structure and function should be further investigated.

### 4.2 Aphids respond to different parasitoid lineages

Aphids in this experiment showed a small shift in gene expression when housing different parasitoid lineages (Analysis 2, Figure 3). We emphasize that this comparison was entirely among *H. defensa*-free aphids from a single clone, hence these differences were not due to the aphid genotype nor its microbiome. That the differences in aphid gene expression were limited is not surprising, considering that this aphid clone is more susceptible to parasitoid attack without *H. defensa* and because parasitoid attack in general will impose many similar stresses (Dennis *et al.* 2017). These differences are potentially informative, though, because they should represent aphid responses to recently evolved differences among the wasps.

Differentially expressed genes in aphids included chitinase, proteins regulating transport to the Golgi bodies, transcription factors, and KEGG pathways associated with metabolism. Unfortunately, many candidates for aphid defenses could not be identified. One explanation for this is that these are not well-characterized genes. While genome-wide annotation work has identified a number of missing immune pathways, notably including antimicrobial activity (Gerardo *et al.* 2010; Laughton *et al.* 2011), this approach would not identify pathways that are specific to aphids. For this, detailed work would be needed to isolate defenses specific to individual aphid species or clones, as has been done recently to identify novel interactions between symbionts and aphid immune cells (Schmitz *et al.* 2012). While from a very disparate system, previous work in *Drosophila* has also found that their response to parasitoid attack was quite dissimilar to known immune pathways (Wertheim 2015).

### 4.3 Evolved expression differences underlie variation in parasitoid infectivity

The most striking comparison of gene expression comes from parasitoid larvae reared in a common environment (H- aphids, Analysis 2). Among the three treatments, several hundred genes were differentially expressed. These expression differences are expected to contribute to infective differences among parasitoid lineages, and their expression in a common environment suggests that they are genetically based. Among these, we see genes with putative functions towards both toxicity and detoxification, and regulators of gene expression. This is a system filled with toxins, in both the aphids’ defenses and the wasp venom; thus it seems that parasitoid success could be the result of the specificity in these products (Colinet *et al.* 2014; Colinet *et al.* 2013), while simultaneously diffusing incoming toxins from the host (Oliver & Higashi 2018).

Among genes that could help in detoxification are additional ribosomal proteins, which we have discussed above. There are also several heat shock proteins; these key molecular chaperones are widely seen as a first defense in stabilizing proteins under both biotic and abiotic stressors (King & MacRae 2015). We also see the differential expression of cytochrome p450, best known in insects for its role in metabolizing insecticides and other toxins, but also upregulated under temperature stress (Feyereisen 1999; Jedlička *et al.* 2015; Scott Jeffrey & Wen 2001).

Putative toxin genes bear similarity to those previously identified in other parasitoids, from the toxins of other animals, and among the differentially expressed genes in adult females from these lineages (Dennis *et al.* 2017). These include F-box leucine-rich repeats, metalloproteinases, and venom carboxylesterases. In adult females more putative venom toxins were differentially expressed than in the larvae (Supplemental data 9, Dennis *et al.* 2017), and it seems unlikely that larvae at this stage are producing venom. Thus, we suggest that genes with this function that are expressed in larvae could instead be secreted, and function to compromise aphid reproduction and defenses. This has been demonstrated in a *Drosophila* parasitoid (*Microplitis mediator*), where production of a metalloproteases disrupts the host Toll pathways; we see this immune pathway upregulated in infected aphids (Lin *et al.* 2018). For the most part, genes with this function were more highly expressed in the H- and H402 treatments. The H76 lineage of wasps may overexpress toxins that we did not identify, or express these at different developmental time points (Martinez *et al.* 2016).

The strongest candidates for evolved expression differences among treatments come from genes that are differentially expressed in both adults and larvae (Figure 5). This includes proteins associated with detoxification and protein stability, namely heat shock and ribosomal proteins, as discussed above. Importantly, a leucine-rich-repeat protein is repeatedly differentially expressed, suggesting that manipulation of this venom component in particular could be important. We also see repeated occurrence of *Nedd4*-binding protein 2; three genes matching to this are upregulated in both adults and larvae of the H402 treatment only (Figure 5). While their role here is not clear, these proteins bind to the highly conserved *Nedd4*, an E3 ubiquitin ligase that has been localized in developing neural tissue in mammals (Kumar *et al.* 1997). *Nedd4* activity has also been associated with promotion of replication by Japanese encephalitis, a ssRNA(+) virus, in humans (Xu *et al.* 2017), so it is possible that these parasitoid lineages differ in their viral susceptibility. This is interesting both because of the very high viral load observed in these samples, and previous identification of a ssRNA(+) virus with lower presence in adult females from the H402 lineage (Dennis *et al.* 2017; Lüthi *et al.* in prep).

### 4.4 Plastic response to the presence of *H. defensa* is small in parasitoids

We examined plastic responses of parasitoid larvae to the presence of *H. defensa* by comparing just one treatment (H76 or H402) when reared in aphids that did or did not possess the *H. defensa* strain with which the wasps had evolved. In the H76 treatment, we can see a further upregulation of candidate genes including leucine-rich-repeat proteins, and ribosomal proteins (L4) when the aphids carry *H. defensa* strain H76, suggesting plasticity in the scope of their expression. However, we do not see such patterns in the H402 treatment. It is important to note that increased infectivity of the H76 and H402 treatments only occurs in the presence of *H. defensa*, meaning that they do not have measurably increased fitness in the control (H-) conditions (Dennis *et al.* 2017). Therefore, we suggest that these small, plastic shifts in gene expression are in direct response to the toxins produced by *H. defensa*, and do not counteract any defenses produced by the aphids themselves.

### 4.5 Viral load

Unexpectedly, some samples contained a large proportion of putatively viral reads. We excluded these from all analyses, but their presence is nonetheless noteworthy and warrants future investigation. Viruses have been identified from a growing number of RNA-seq studies, including insects (Liu *et al.* 2015; Obbard 2018; Shi *et al.* 2016; Webster *et al.* 2015), and parasitoids in particular (Oliveira *et al.* 2010; Renault 2012). We have previously identified an abundant ssRNA (+) virus, provisionally named LysV, in both these experimental evolution lineages and in the wild (Dennis *et al.* 2017; Lüthi *et al.* in prep), and this virus was also present in the current study. However, many of the putatively viral sequences found here appear to belong to another lineage of RNA-viruses and to a newly described lineage thus far called “dark matter” viruses (Obbard 2018). As with LysV, we believe the abundant viral sequences identified here are from *L. fabarum*, and not their aphid hosts because they: (a) are only abundant in libraries where wasps were understood to be healthy, (b) were not detected in another RNAseq study of the same aphid lines uninfected by parasitoids (H. Käch, *personal communication*), and (c) were present in the libraries generated from adult females. We have not observed any indication that these viruses have impacted parasitoid fitness, although this should be further investigated. Given their persistence in the lab for >15 generations, it is possible that there could be some benefit to hosting these viruses, as has been observed in other parasitoids (Dheilly *et al.* 2015a; Dheilly *et al.* 2015b; Martinez *et al.* 2012).

### 4.5 Conclusion

Here we have used dual RNA-seq to examine functional variation between experimentally evolved parasitoid populations and their aphid hosts. Despite their strong reliance on the endosymbiont *H. defensa* for protection, subtle changes in aphid gene expression suggest that aphids are not passive bystanders. In parasitoids, we have identified differences in expression of genes that include detoxifying and putative venom components. Importantly these differences appear to be evolved rather than plastic upregulation in the presence of *H. defensa*, since they were also observed when *H. defensa*-adapted parasitoids were reared in *H. defensa*-free aphids. Together, this paints a picture of a variable parasitoid strategy to cope with the defenses brought on by the aphid microbiome, providing strong candidates for the direct mechanisms by which wild parasitoid populations maintain diverse strategies to cope with the symbiont-mediated defenses of their hosts.

## 5 Acknowledgements

Special thanks to Hannele Penson and Paula Rodriguez for assistance with insect rearing, to Martina Lüthi and Pravin Ganesanandamoorthy for help with many double RNA-DNA extractions, and Stefanie Hartmann for bioinformatics assistance. Data produced and analyzed in this paper were generated in collaboration with the Genetic Diversity Centre (GDC), ETH Zurich. This study was supported by the Swiss National Science Foundation (SNSF Professorship nr. PP00P3_146341 and Sinergia grant nr. CRSII3_154396 to CV) and personnel support from EAWAG.

## 7. Data accessibility

Illumina data has been deposited in NCBI short read archives (SAMN10024115-SAMN10024165). The *de novo* assemblies for both the parasitoid *L. fabarum* and host aphid *A. fabae* will be deposited in Dryad, along tables of the top *blast*-hit of each assembled contig (*doi pending manuscript publication, happy to provide this data in the meantime if needed*).

## 8. Author Contributions

CV conceived of the experimental evolution design and carried it out with ABD. ABD designed and carried out differential expression analyses. HK provided transcriptomic sequences from aphids. All three authors contributed to writing this manuscript.

## 11. Supporting information

List of supplementary data files/ results:

1. Analysis 1, DGE in aphids. Overall analysis of differential expression in aphids-comparing infected to uninfected individuals
2. Analysis 1, KEGG analysis in aphids. KEGG analysis of expression between infected and uninfected aphids
3. Analysis 1, DGE in parasitoids. Overall analysis of differential expression in parasitoid wasp larvae-comparing successful to unsuccessful individuals
4. Analysis 1, KEGG analysis in parasitoids. KEGG analysis of expression between successful and unsuccessful parasitoid wasps
5. Analysis 2, DGE among only H- aphids, hosting three parasitoid treatments.
6. Analysis 2, KEGG analysis among only H- aphids, hosting three parasitoid treatments.
7. Analysis 2, DGE among larval parasitoids from three treatments, only hosted in H- aphids.
8. Analysis 2, KEGG analysis among larval parasitoids from three treatments, only hosted in H- aphids.
9. Analysis 2, DGE among adult parasitoids from three treatments, only hosted in H- aphids.
10. Analysis 2, KEGG analysis among adult parasitoids from three treatments, only hosted in H- aphids.
11. Analysis 3 (treatment 402): DGE of parasitoid from the H402 treatment wasps housed by either H- or H402 aphids
12. Analysis 3 (treatment 76): DGE of parasitoid from the H76 treatment wasps housed by either H- or H76 aphids

**Supplemental Table 1:** Summary of all dual RNA-seq libraries, divided into number of reads mapped to the *L. fabarum* and *A. fabae* reference transcriptomes, and to the six putative-viral transcripts identified in the data. Viral reads were not included in any analyses.

| Sample number | Successful parasitoid infection? | Parasitoid treatment | Host aphid | WASP mapped reads | APHID mapped reads | Viral read count | Notes |
| --- | --- | --- | --- | --- | --- | --- | --- |
| 100_1 | yes | 402.1 | H- | 3,938,329 | 12,774,395 | 37,172 | § |
| 100_2 | yes | 402.1 | H- | 1,499,082 | 16,257,704 | 35,425 |  |
| 103_19 | yes | 402.2 | H- | 9,641,147 | 5,483,491 | 46,862 |  |
| 104_23 | yes | 402.3 | H- | 987,204 | 629,396 | 14,106,378 | †‡ |
| 104_24 | yes | 402.3 | H- | 1,506,538 | 2,561,305 | 10,284,530 |  |
| 105_34 | yes | 402.3 | H- | 2,869,983 | 3,571,536 | 10,055,675 |  |
| 107_48 | yes | 402.4 | H- | 13,254,955 | 2,262,943 | 66,682 |  |
| 107_51 | yes | 402.4 | H- | 12,554,226 | 1,292,525 | 67,300 | ‡ |
| 107_52 | yes | 402.4 | H- | 168,595 | 33,240 | 156,167 | †‡ |
| 110_360 | no | 402.2 | 402 | 1,795,348 | 16,204,372 | 242,762 | ¶ |
| 110_363 | yes | 402.2 | 402 | 10,815,918 | 3,611,004 | 140,777 |  |
| 110_365 | no | 402.2 | 402 | 1,749,097 | 11,900,248 | 189,038 | ¶ |
| 113_446 | yes | 402.3 | 402 | 520,458 | 6,244,051 | 10,095,306 | † |
| 113_448 | yes | 402.3 | 402 | 1,867,533 | 5,226,079 | 7,852,322 |  |
| 113_449 | yes | 402.3 | 402 | 2,395,408 | 7,976,771 | 7,171,124 |  |
| 114_483 | yes | 402.4 | 402 | 32,347 | 7,288 | 109,613 | †‡ |
| 114_488 | yes | 402.4 | 402 | 4,948,264 | 1,000,239 | 42,840,636 | ‡ |
| 114_491 | yes | 402.4 | 402 | 5,712,357 | 2,158,694 | 115,706 |  |
| 115_519 | no | 402.4 | 402 | 2,355,958 | 19,631,427 | 241,771 | Ø |
| 115_520 | no | 402.4 | 402 | 2,807,946 | 22,400,244 | 322,724 | Ø |
| 115_522 | no | 402.4 | 402 | 4,191,451 | 9,849,741 | 393,691 | Ø |
| 132_568 | no | 76.1 | 76 | 1,881,205 | 18,809,189 | 199,163 | ¶ |
| 132_569 | no | 76.1 | 76 | 3,915,620 | 10,623,689 | 295,852 | ¶ |
| 135_646 | yes | 76.2 | 76 | 268,674 | 254,986 | 10,904,890 | †‡ |
| 137_684 | yes | 76.3 | 76 | 746,956 | 4,637,224 | 8,618,935 | †‡ |
| 137_686 | no | 76.3 | 76 | 58,995 | 1,017,572 | 5,898 | † Ø |
| 137_693 | no | 76.3 | 76 | 1,045,578 | 4,012,416 | 65,967 | Ø |
| 137_696 | no | 76.3 | 76 | 193,671 | 3,294,192 | 18,665 | † Ø |
| 137_699 | yes | 76.3 | 76 | 571,738 | 8,187,020 | 7,868,131 | †‡ |
| 124_202 | yes | 76.1 | H- | 2,314,967 | 459,297 | 13,212,453 | ‡ |
| 124_203 | yes | 76.1 | H- | 9,733,902 | 1,751,539 | 9,075,656 |  |
| 126_225 | yes | 76.2 | H- | 3,482,944 | 2,508,033 | 11,858,633 |  |
| 126_227 | yes | 76.2 | H- | 1,454,623 | 558,340 | 10,792,907 | ‡ |
| 126_228 | yes | 76.2 | H- | 2,056,255 | 511,932 | 13,002,679 | †‡ |
| 129_268 | no | 76.3 | H- | 369,521 | 9,333,442 | 6,812,122 | †¶ |
| 130_269 | yes | 76.4 | H- | 3,898,792 | 4,466,715 | 10,536,556 |  |
| 130_273 | yes | 76.4 | H- | 5,762,995 | 3,961,447 | 12,839,652 |  |
| 145_89 | yes | 407.1 | H- | 19,983,577 | 27,941,295 | 128,071 |  |
| 145_90 | yes | 407.1 | H- | 55,594,294 | 18,534,512 | 476,165 |  |
| 145_91 | yes | 407.1 | H- | 4,656,974 | 4,070,798 | 79,773 |  |
| 118_111 | yes | 407.2 | H- | 2,544,011 | 2,081,936 | 4,163,964 |  |
| 118_116 | yes | 407.2 | H- | 3,019,968 | 2,730,510 | 840,937 |  |
| 118_118 | yes | 407.2 | H- | 6,816,139 | 10,990,187 | 53,132,336 |  |
| 142_129 | no | 407.2 | H- | 2,110,279 | 24,078,429 | 131,093 | Ø |
| 142_131 | no | 407.2 | H- | 1,982,643 | 14,865,386 | 121,670 | Ø |
| 142_132 | no | 407.2 | H- | 2,430,970 | 24,577,577 | 264,575 | Ø |
| 120_145 | yes | 407.3 | H- | 57,337,440 | 21,446,654 | 41,642,000 |  |
| 120_147 | yes | 407.3 | H- | 31,683,727 | 15,936,026 | 100,385,598 |  |
| 120_151 | yes | 407.3 | H- | 21,806,153 | 18,211,786 | 58,935,883 |  |
| 122_174 | no | 407.4 | H- | 4,480,101 | 43,349,507 | 453,494 | ¶ |
| 141_196 | no | 407.4 | H- | 113,782 | 2,960,102 | 19,758 | †¶ |

† dropped due to low read-count (parasitoid)
‡ dropped due to low read-count (aphid)
§ outlier, removed from all analyses
¶ designaged uninfected by expression profile
Ø aphid not detected by PCR, deemed uninfected

**Supplemental Table 2:** Summary of the six abundant transcripts that are likely viruses in the system

| Contig name | AA length | Mapped reads | Top hit (blasp) | e-value | Accession of match | Category of match |
| --- | --- | --- | --- | --- | --- | --- |
| c321_g1_i1 | 574 | 4,589,238 | capsid [Culex pipiens associated Tunisia virus] | 2.00E-27 | AUT77207 | RNA virus |
| c25231_g1_i1 | 265 | 36,909,654 | hypothetical protein [Kwi virus] | 5.00E-40 | AVB77240 | unclassified viruses |
| c29007_g2_i1 | 599 | 204,138,574 | hypothetical protein [Nai virus] | 1.00E-45 | AVB77237 | unclassified viruses |
| c29502_g1_i1 | 584 | 81,371,681 | hypothetical protein [Kwi virus] | 5.00E-53 | AVB77242 | unclassified viruses |
| C29860_g1_i1 | 728 | 57,013,829 | putative RNA polymerase [Kwi virus] | 0 | AYU49214 | unclassified viruses |
| c31332_g1_i1 | 608 | 96,662,281 | hypothetical protein [Nai virus] | 4.00E-102 | AVB77235 | unclassified viruses |

